# Integrating Vectors for Genetic Studies in the Rare Actinomycete *Amycolatopsis marina*

**DOI:** 10.1101/436022

**Authors:** Hong Gao, Buvani Murugesan, Janina Hoßbach, Stephanie K. Evans, W. Marshall Stark, Margaret C.M. Smith

## Abstract

Few natural product pathways from rare Actinomycetes have been studied due to the difficulty in applying molecular approaches in these genetically intractable organisms. In this study, we sought to identify integrating vectors, derived using phage *int/attP* loci, that would efficiently integrate site-specifically in the rare Actinomycete, *Amycolatopsis marina* DSM45569. Analysis of the genome of A. *marina* DSM45569 indicated the presence of *attB*-like sequences for TG1 and R4 integrases. The TG1 and R4 *attBs* were active in *in vitro* recombination assays with their cognate purified integrases and *attP* loci. Integrating vectors containing either the TG1 or R4 *int/attP* loci yielded exconjugants in conjugation assays from *E. coli* to *A. marina* DSM45569. Site-specific recombination of the plasmids into the host TG1 or R4 *attB* sites was confirmed by sequencing. The presence of homologous TG1 and R4 *attB* sites in other species of this genus indicates that vectors based on TG1 and R4 integrases could be widely applicable.

**Importance:** Rare Actinomycetes have the same potential of natural product discovery as Streptomyces, but the potential has not been fully explored due to the lack of efficient molecular biology tools. In this study, we identified two serine integrases, TG1 and R4, which could be used in the rare Actinomycetes species, *Amycolatopsis marina*, as tools for genome integration. The high level of conservation between the *attB* sites for TG1 and R4 in a number of Amycolatopsis species suggested that plasmids with the integration systems from these phages should be widely useful in this genus.

## Introduction

*Streptomyces* bacteria are widely exploited for their abundant bioactive natural products(1). However, after 70 years of exploitation, the rate of discovery of new *Streptomyces-de*rived bioactive products has declined, and interest in other potential non-Streptomycete sources, such as the rare Actinomycetes, has grown(2, 3).

Amongst rare Actinomycetes, the *Amycolatopsis* genus is of particular interest for its production of important antibiotics such as vancomycin(4) and rifamycin(5), as well as a diverse range of active natural products(6-8). The publicly available NCBI database contains nearly 70 genomes of *Amycolatopsis* strains, covering more than 40 species from this genus. Similar to *Streptomyces*, the genome of each *Amycolatopsis* contains, on average, over 20 secondary metabolic gene clusters(9). The mining of these metabolic clusters offers great potential for novel antibiotic discovery. However, the lack of widely available and efficient genetic tools for these rare species impedes their potential in research and development.

Phage-encoded serine and tyrosine integrases catalyse site-specific integration of a circularized phage genome into the host chromosome as part of the process to establish a lysogen. DNA integration mediated by serine integrases occurs between short (approximately 50 bp) DNA substrates that are located on the phage genome, (the phage attachment site *attP*) and the host genome (the bacterial attachment site *attB*). The product of *attP* x *attB* recombination is an integrated phage genome flanked by two new sites, *attL* and *attR*, each of which contains half-sites from *attP* and *attB*. During phage induction, integrase in the presence of a recombination directionality factor (RDF) again mediates site-specific recombination, but this time between *attL* and *attR*, to excise the phage genome, which can then be replicated during a lytic cycle. The mechanism of recombination and the factors that control integration versus excision have been elucidated in recent years(10-12).

Integrating vectors based on the *Streptomyces* phage ϕC31 integrase and *attP* locus are the best known and most widely used in Actinomycete genome engineering(13-16). In addition to the phage recombination machinery (*int/attP*), integrating vectors contain a replicon for maintenance in *E. coli*, an *oriT* for conjugal transfer and a marker or markers for selection in *E. coli* and in the recipient. They are powerful genome engineering tools that act in an efficient, highly controllable and predictable way(17).

Integrating vectors using serine integrase-mediated recombination require no additional phage or host functions for integration, an especially important feature when they are to be used in other organisms that cannot be infected by the phages. This property makes serine integrase-based vectors promising tools for use in heterologous systems(11, 18). However, use of these integration vectors has not been fully explored in rare Actinomycetes, e.g. *Amycolatopsis*. There is only one reported example of a conjugation system based on ϕC31 integrase in *Amycolatopsis Japonicum* MG4I7-CFI7(19), and it has been reported that other *Amycolatopsis* species lack ϕC31 *attB* sites in their chromosomes(20). The ϕBT1 *attB* sites have been more commonly identified in *Amycolatopsis*, and there has been successful conjugative transfer of vectors based on ϕBT1 *int/attP* in *A. orientalis(20)* and *A. mediterranei*(21). Furthermore, electroporation remains the most widely applied method for transfer of integrative plasmids into this genus, rather than conjugation(20, 21).

In this paper, we chose to study *Amycolatopsis marina* DSM45569, a species isolated from an ocean-sediment sample collected in the South China Sea(22). We explored the application of bacterial genetic engineering using serine integrases, and developed conjugative and integrating vectors for use in this species. We present evidence suggesting that these vectors could be applied to various other species in this genus, opening up the prospect for more versatile genetic manipulation of *Amycolatopsis*.

## Materials and Methods

### Bacterial strains and culture conditions

Plasmid propagation and subcloning was conducted using *E. coli* Top10 (F- *mcrA* Δ(*mrr-hsdRMS-mcrBC*) ϕ80*lac*ZΔM15 *ΔlacX74 nupG recA1 araD139* Δ(*ara-leu)7697 galE15 galK16 rpsL*(Str^R^) *endA1* λ^-^). Plasmid conjugations from *E. coli* to *A. marina* DSM45569 were carried out using *E. coli* ET12567(pUZ8002) containing the plasmid to be transferred as the donor (23, 24), and conjugations from *E. coli* to *S. coelicolor* and *S. lividans* were used as control. *E. coli* strains were grown in Luria-Bertani broth (LB) or on LB agar at 37°C.

*Amycolatopsis marina* DSM45569 was purchased from the German Collection of Microorganisms and Cell Cultures (DSMZ, Germany), and maintained on Soya Mannitol (SM) agar at 30°C. Harvested spores were maintained long-term in 20% glycerol at −80°C. Conjugations were plated on SM agar containing 10 mM MgCl_2_, and ISP2 medium(25) was used for the preparation of genomic DNA(24).

### DNA manipulation

*E. coli* transformation and gel electrophoresis were carried out as described previously(26). Genomic DNA preparation from *Streptomyces* was performed following the salting out procedure in the *Streptomyces* manual(24). Plasmids from *E. coli* were prepared using QIAprep^®^ Spin Miniprep Kit (Qiagen, Germany) following the manufacturer’s instructions. Polymerase Chain Reaction (PCR) was carried out using Phusion^®^ High-Fidelity DNA Polymerase (NEB, USA) according to the manufacturer’s instructions. The primers used in this study are listed in Table 1. DNA samples were purified by the QIAquick Gel Extraction Kit (Qiagen, Germany).

**TABLE 1.**
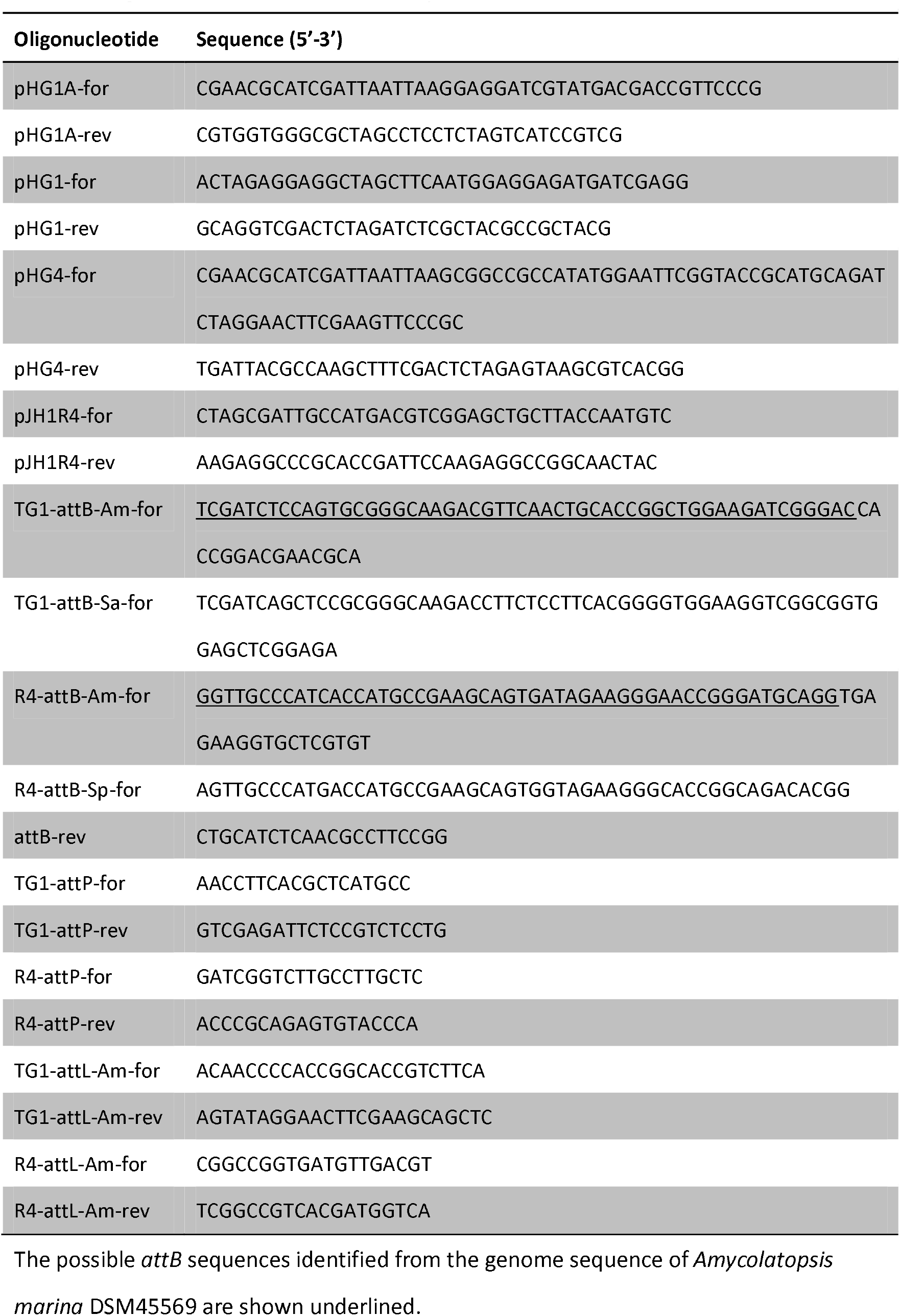
Oligonucleotides used in this study

### Plasmid construction

The integrating plasmid pHG4 contains the TG1 *int*/*attP* locus and the apramycin-resistance gene (*aac3*(*IV*)) for selection (Figure 1A). The fragment containing *oriT, aac3*(*IV*) and TG1 *int/attP* was amplified from plasmid pBF20(27) using the primer pair pHG4-for/pHG4-rev. The fragment was joined via In-Fusion cloning to the 3344 bp HindIII-Pacl fragment from pBF22(27) (containing the *E. coli* plasmid replication origin, the *bla* gene encoding resistance to ampicillin and the *actll-orf4*/*act1p* expression cassette) to form the plasmid pHG4.

**Figure 1.**
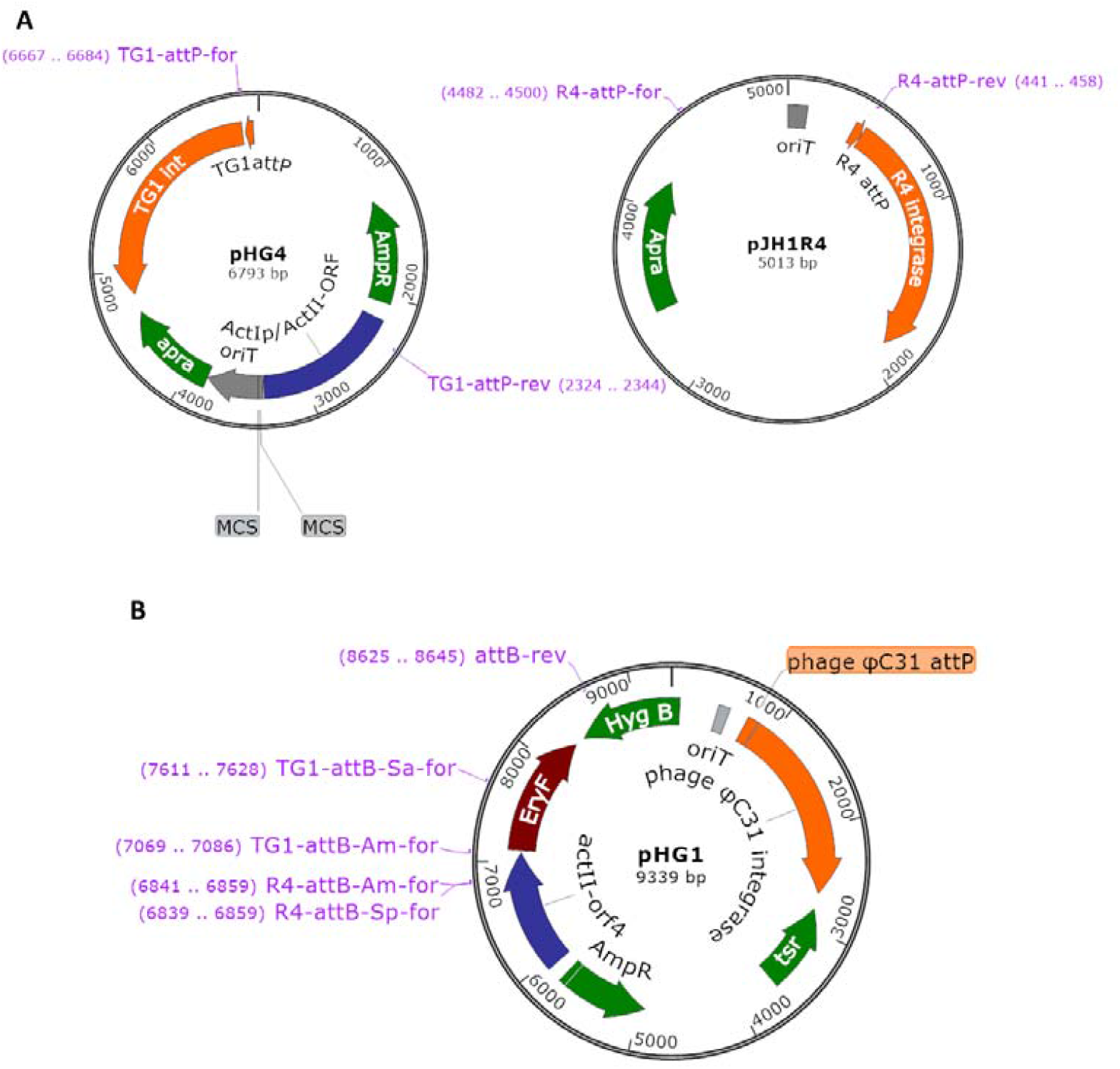
Plasmids used in this study. The primer binding sites are indicated. (A) Integrating plasmids pHG4 and pJH1R4; (B) PCR template plasmid pHG1.

To construct the integrating plasmid pJH1R4 (Figure 1A), pSET152(28) was cut with Aatll and Pvul to remove the ϕC31 *attP* site and integrase gene. R4 phage lysate was used as template in a PCR with the primers pJH1R4-for and pJH1R4-rev to amplify the R4 *attP* site and integrase coding region. The PCR product was joined to the Aatll-Pvul fragment from pSET152 via In-Fusion cloning.

The plasmid pHG1 (Figure 1B) was used as template to amplify *attB*-containing sequences for *in vitro* recombination assays. This plasmid was originally constructed for the expression of *EryF*. The *eryF* gene was amplified from *Saccharopolyspora erythraea* BIOT-0666 genomic DNA using the primer pair pHG1A-for/pHG1A-rev, and inserted by In-Fusion cloning into pBF20(27) cut with Nhel and Pacl to form the plasmid pHG1A. The 3785 bp fragment containing the ϕC31 *int/attP* and hygromycin resistance gene was amplified from plasmid pBF27C(27), using the primer pair pHG1-for and pHG1-rev. Plasmid pHG1A was digested with Xbal and Nhel, and the 5668 bp fragment was ligated with the 3785 bp PCR fragment from pBF27C by In-Fusion to give the plasmid pHG1.

### *In Vitro* Recombination Assays

*In vitro* recombination assays were performed using PCR-amplified DNA fragments containing the *attB* and *attP* attachment sites located at the ends. Recombination between the *attP* and *attB* sites joined the two fragments to give a product whose length was almost the sum of the substrates (Figure 3A). To generate the *attB*-containing substrates, the forward primer, TG1-attB-Am-for, contained the closest match in the A. *marina* genome to the characterised TG1 *attB* site from *S. avermitilis*, TG1 *attB*^Sa^ (29)(Figure 2). TG1-attB-Am-for also had a sequence identical to the 3’ end of the act1p element from plasmid pHG1, which was used as a template for PCR (Figure 1). Similarly, the forward primer R4-attB-Am-for contained the closest match in the A. *marina* genome to the characterized R4 *attB* site from *S. parvulus*, R4 *attB^Sp^* (30) (Figure 2). R4-attB-Am-for also had a sequence identical to the 3’ end of Actll-orf4 element from the template plasmid pHG1 (Figure 1). Forward primers TG1-attB-Sa-for and R4-attB-Sp-for were used to create positive control recombination substrates containing the TG1 and R4 *attB* sites originally found in S. *avermitilis*(29) and *S. parvulus*(30) respectively. The reverse primer used to generate all the *attB*-containing substrates (attB-rev) was located within the *hyg* gene of pHG1; the amplified products were 1627 bp (TG1 *attB*^Am^), 1035 bp (TG1 *attB*^Sa^), 1854 bp (R4 *attB*^Am^) and 1855 bp (R4 *attB^Sp^*). The DNA fragments containing the *attP* sites were prepared as follows; the TG1-*attP* fragment (2471 bp) was amplified using the primer pair TG1-attP-for/TG1-attP-rev with pHG4 as the template, and the R4-*attP* fragment (990 bp) was amplified using the primer pair R4-attP-for/R4-attP-rev with pJH1R4 as the template (Figure 1). Note that other than the *attB* and *attP* sites, none of the substrates contained any DNA that should interact specifically with the integrases. Moreover, each fragment was designed to be easily identifiable by molecular weight. The integrases were purified as described previously(31, 32). All recombination reactions were in 20 μl final volume. Recombination reactions of TG1 substrates were carried out in TG1 RxE buffer (20 mM Tris [pH 7.5], 25 mM NaCl, 1 mM dithiothreitol [DTT], 10 mM spermidine, 10 mM EDTA, 0.1 mg/ml bovine serum albumin [BSA])(33), and recombination reactions of R4 substrates were carried out in buffer containing 20 mM Tris-HCl (pH 7.5), 50 mM NaCl, 10 mM spermidine, 5 mM CaCl_2_ and 50 mM DTT(32). Integrase was added at the concentrations indicated. Recombination substrates were used at 50 ng each per reaction. Reactions were incubated at 30°C overnight, and then heated (10 min, 75°C) to denature integrase. The reaction mixtures were loaded on a 0.8% agarose gel in Tris/Borate/EDTA (TBE) buffer (90 mM Tris base, 90 mM boric acid and 2 mM EDTA) containing ethidium bromide for electrophoretic separation.

**Figure 2.**
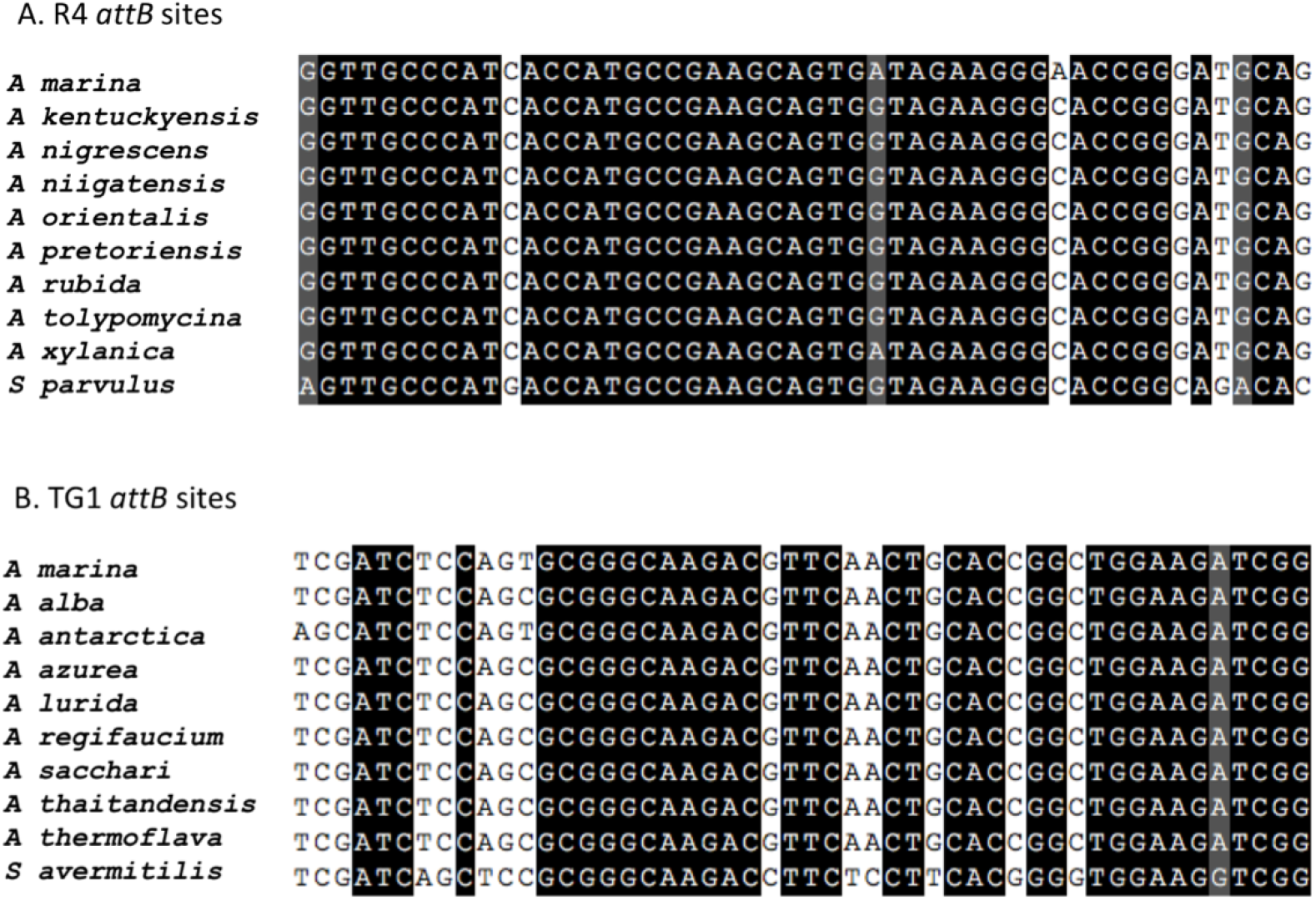
Alignment of R4 and TG1 *attB* sites in *A. marina* DSM45569 and other *Amycolatopsis* species. GenBank accession nos. of DNA sequences: *A. alba* (NMQU01000019.1), *A. antarctica* (NKYE01000021.1), *A. azurea* (MUXN01000005.1), *A. kentuckyensis* (MUMI01000226.1), *A. lurida* (FNTA01000004.1), *A. nigrescens* (ARVW01000001.1), *A. niigatensis* (PJMYO1000003.1), *A. orientalis* (ASXH01000007.1), *A. pretoriensis* (MUMK01000092.1), *A. regifaucium* (LQCI01000034.1), *A. rubida* (FOWC01000001.1), *A. sacchari* (FORP0I0000I0.1), *A. thailandensis* (NMQ.T01000219.1), *A. thermoflava* (AXBH01000004.1), *A. tolypomycina* (FNS001000004.1), *A. xylanica* (FNON01000002.1), *Streptomyces avermitilis* (NC_003155.5) and *S. parvulus* (CP015866.1).

**Figure 3.**
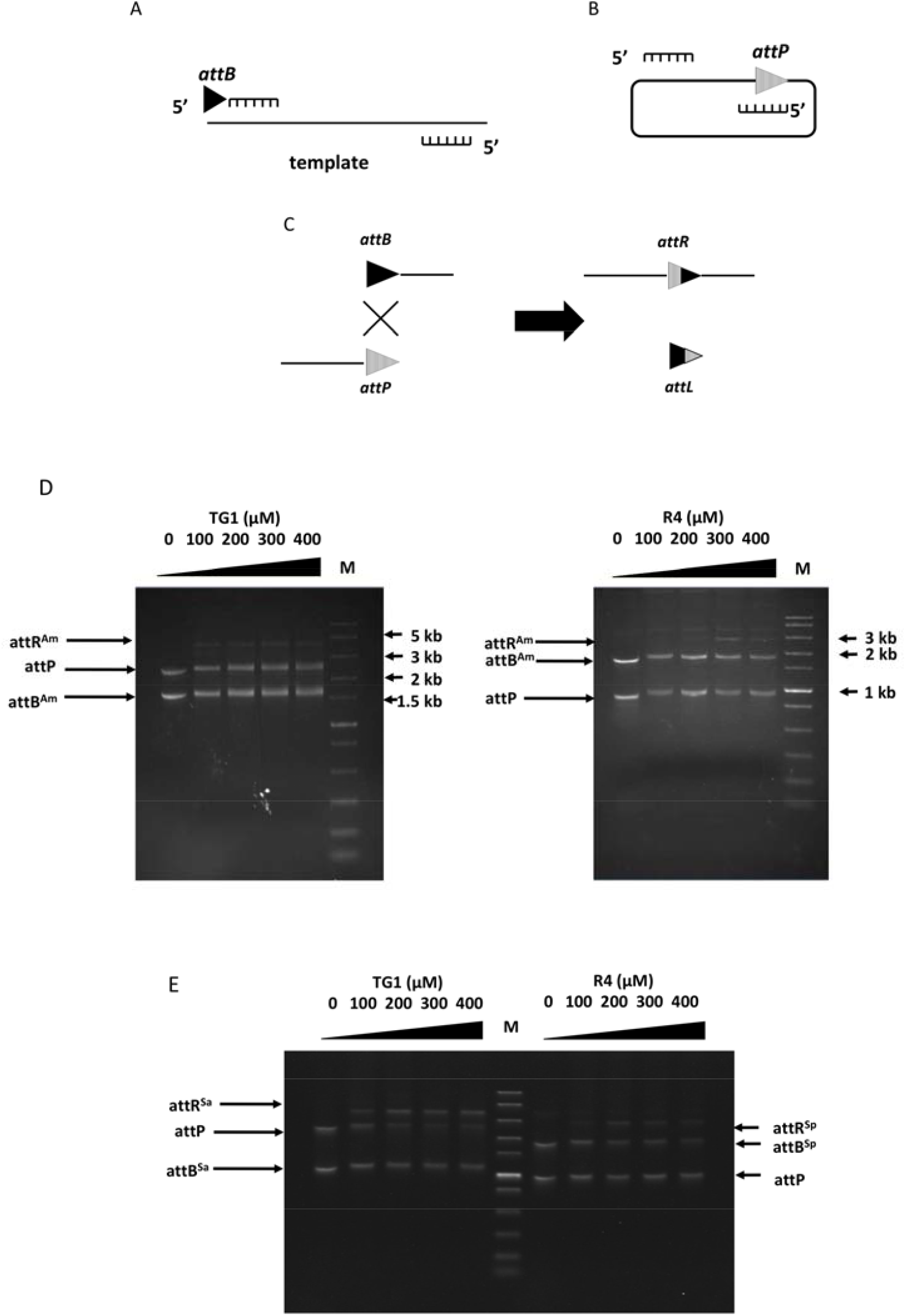
(A) Design and synthesis of DNA substrate *attB*. The *attB* sites were fused at the end of forward primers, to amplify a sequence flanked with *attB* from an unrelated DNA template, pHG1 in this case. (B) Design and synthesis of DNA substrate *attP*. The *attP* sites were amplified directly from integrating vectors carrying TG1/*attP* (pHG4) or R4/*attP* (pJH1R4). (C) Recombination substrates and their expected products. (D) *In vitro* recombination between DNA fragments containing TG1 *attB^Am^*(l627 bp) and TG1 *attP* (2471 bp; left), and R4 *attB^Am^* (1854 bp) and R4*attP* (990 bp; right). The expected products of the TG1 integrase-mediated reaction were a 4.1 kb DNA fragment containing the *attR^Am^* site, and a 53 bp fragment containing *attL^Am^* (not observed). For the R4 integrase recombination reaction, the expected products were a 2.8 kb fragment containing *attR^m^*, and a 51 bp *attL^Am^* fragment (not observed). (E) *In vitro* recombination between DNA fragments containing TG1 *attB^Sa^*(1035 bp) and TG1 *attP* (2471 bp; left), and R4 *attB^Sp^* (1855 bp) and R4 *attP* (990 bp; right). The expected products were a 3.5 kb fragment containing *attR^Sa^* for the TG1 reaction, and a 2.8 kb fragment containing *attR^Sp^*for the R4 reaction. M: Fast DNA Ladder (NEB, USA).

## Results

### Identification of possible *attB* sequences from the genome of *Amycolatopsis marina* DSM45569

The sequences of *attB* sites recognised by a variety of integrases (ϕC31(34), ϕJoe(35), Bxbl(36), R4(32), SPBc(37), SV1(38), TG1(29) and TP901(39)) were used in BLAST searches of the genome sequence of *Amycolatopsis marina* DSM45569 (NCBI Genome Database NZ_FOKG00000000) (Table 2). The most significant hits for R4 and TG1 *attB* sites had the highest identities and lowest E-value. The predicted R4 *attB* site is located within a gene predicted to encode a fatty-acyl-CoA synthase (SFB62308.1) and the TG1 *attB* site is located within a gene predicted to encode a putative succinyldiaminopimelate transaminase (WP_091671332.1). The BLAST analysis was extended to other species of *Amycolatopsis* to assess the conservation of these *attB* sites in the genus (Figure 2). Both R4 and TG1 *attB* sites were highly conserved relative to the *attB* sites originally identified from *S. parvulus*(30) and *S. avermitilis*(29) (84% for R4 and 62% for TG1).

**TABLE 2.**
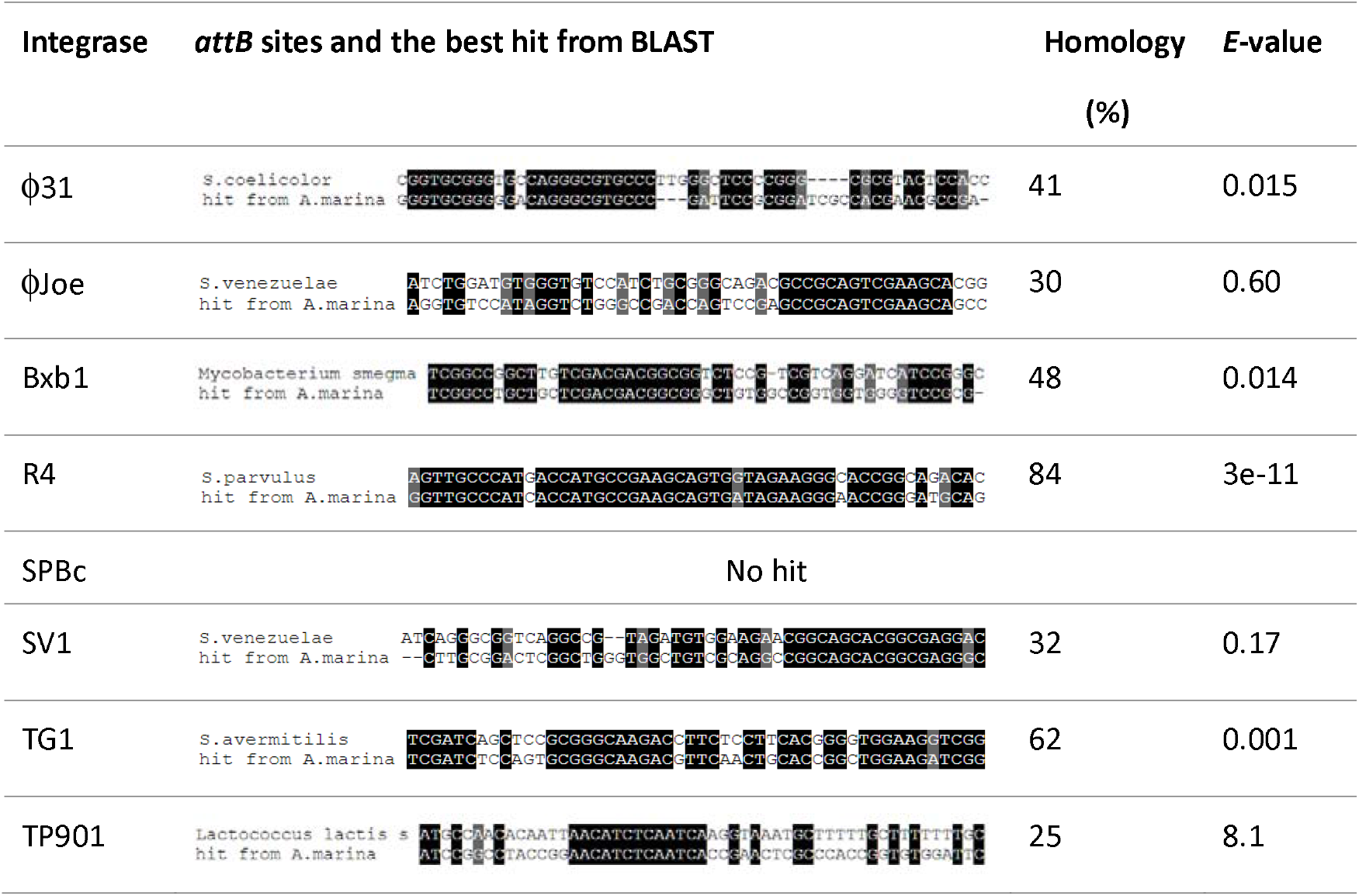
The original *attB* sites for integrases and results of BLAST search

### *A. marinum attB-*1ike sequences for TG1 and R4 are both active in *in vitro* recombination

In each recombination reaction, substrates containing *attP* and the putative *attB* site were mixed in cognate pairs with different concentrations of purified R4 or TG1 integrase in the corresponding buffer and incubated overnight at 30°C, as described in Materials and Methods. The expected recombination events and the nature of the products are shown in Figure 3A-3C. TG1 catalysed recombination between the substrates more efficiently than R4 (Figure 3D). As expected because neither phage is an *Amycolatopsis* phage, the recombination efficiencies for each integrase were observably better when the *Streptomyces attB* sites were used (Figure 3E) compared to the *A. marina attB* sites (Figure 3D), particularly for TG1 integrase. Nevertheless, the presence of recombination activity indicated that both *A. marina attB* sites were functional and were likely to be active integration sites for integrative conjugation vectors.

### *In vivo* integration

*A. marina* DSM45569 was unable to grow in the presence of apramycin, so integrating plasmids pHG4 and pJH1R4, containing the apramycin resistance determinant *aac3(IV)*, were constructed. Following the standard *Streptomyces* conjugation protocol (see Materials and Methods), a frequency of approximately 160 exconjugants/10^8^ spores was obtained for transfer of pHG4 (encoding TG1 integrase), while the conjugation efficiency of pJH1R4 (R4 integrase) was only 20 exconjugants/10^8^ spores (Table 3). For each integration, six exconjugants were picked at random and streaked on MS agar containing apramycin. Genomic DNA was then prepared and used as the template in PCR reactions, in which the primer pairs of TG1-attL-Am-for/rev and R4-attL-Am-for/rev were used to test for the occurrence of recombination at the expected TG1 and R4 *attB* sites (Figure 4). All PCR reactions using exconjugants as templates gave the expected band sizes. Sequencing (GATC, Germany) of the PCR products with the primers TG1-attL-Am-for and R4-attL-Am-for confirmed that the plasmids had integrated into the predicted *attB* sites for TG1 or R4 integrase within *A. marina* DSM45569 (Figure 5).

**Figure 4.**
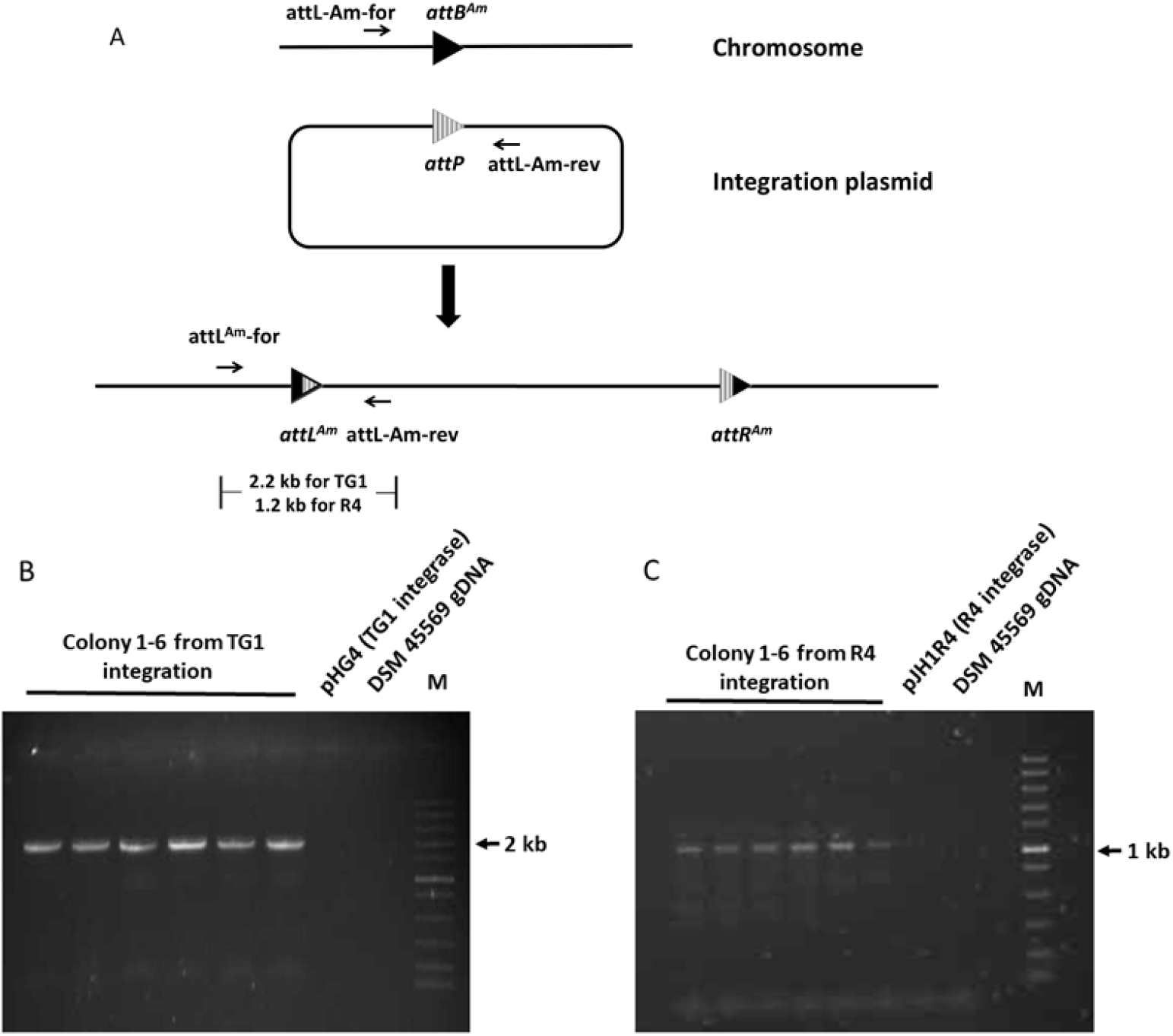
PCR confirmation of site-specific integration in the exconjugants. (A) Design of primers. (B) PCR (using primers TG1-attL-Am-for/rev) of the expected TG1 *attL*-containing fragment from *A. marina* DSM45569:pHG4. (C) PCR (using primers R4-attL-Am-for/rev) of the expected R4 *attL*-containing fragment from *A. marina* DSM45569:pJH1R4. M: Fast DNA Ladder. Colonies 1 to 6 are independent exconjugants.

**Figure 5.**
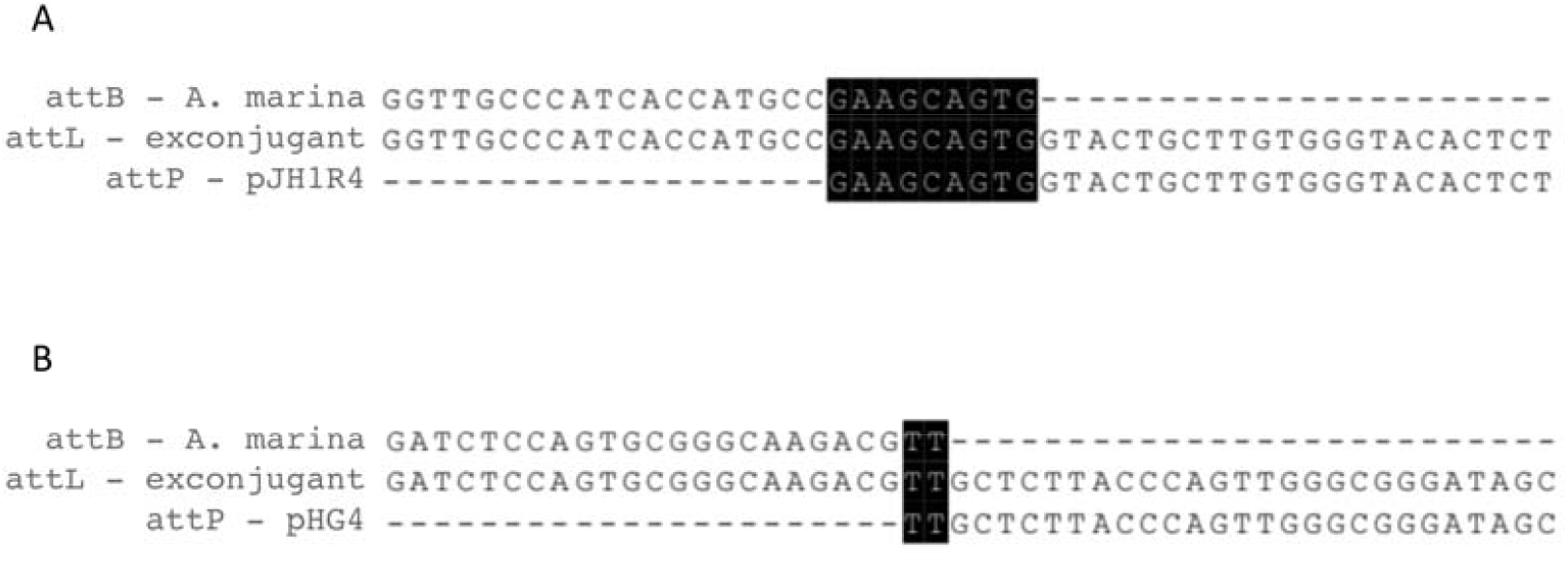
The insertion sites of TG1 and R4 integration plasmids in *A. marina* DSM45569. Sequencing (using primers TG1-attL-Am-for or R4-attL-Am-for) of PCR products containing *attL* from exconjugants validated the site-specific recombination of the TG1 and R4 *attB* sites in *A. marina* DSM45569 after introduction of pHG4 or pJH1R4, respectively.

**TABLE 3.**
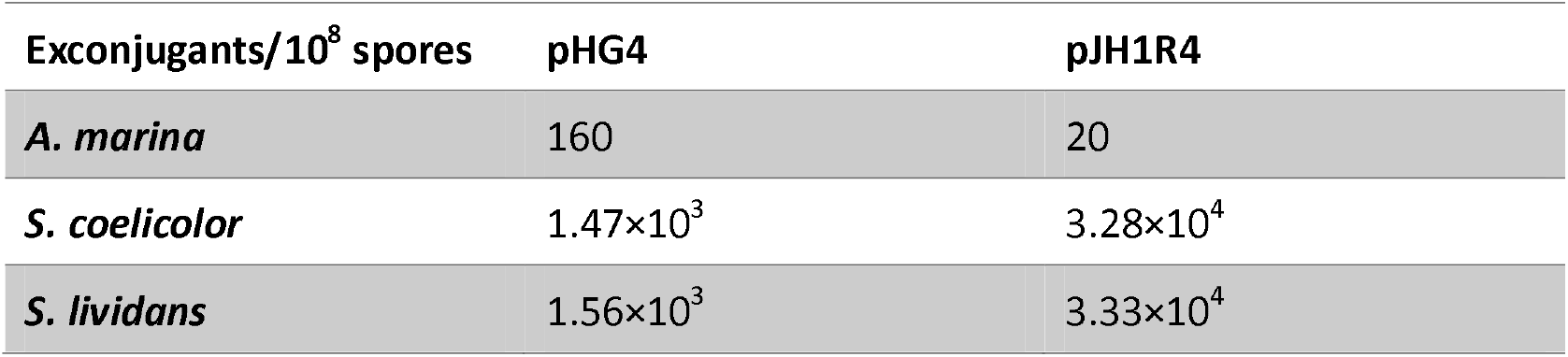
Conjugation efficiency of pHG4 and pJH1R4 in different species

| Exconjugants/10<sup>8</sup> spores | pHG4 | pJH1R4 |
| --- | --- | --- |
| *A. marina* | 160 | 20 |
| *S. coelicolor* | 1.47×10<sup>3</sup> | 3.28×10<sup>4</sup> |
| *S. lividans* | 1.56×10<sup>3</sup> | 3.33×10<sup>4</sup> |

## Discussion

The lack of effective genetic engineering tools is considered one of the greatest hindrances in the search for novel natural products from rare Actinomycetes(40-42). Previous studies in rare Actinomycete species have largely focused on the use of the well-characterised ϕC31-based integration vectors, and have mostly overlooked tools based on other phage integrases(43-45). Additionally, the conjugation methods used widely in *Streptomyces* gene transfer because of their ease have shown little success in rare Actinomycetes, including species in the genus *Amycolatopsis*, so electroporation has been the long-preferred method of gene transfer for species in this genus(5, 9, 46, 47). However, the growing interest in the use of serine integrases for synthetic biology applications(11) has led to further research into expanding the pool of available enzymes and their potentials as genetic tools(48-50). Therefore, within this study, we explored whether integrating vectors based on eight different serine integrases could be employed for genetic engineering of A. *marina* DSM45569. Sequence analysis of the A. *marina* DSM45569 genome identified close matches to the *attB* sites used by TG1 and R4 integrases. Although conjugation frequencies were low, integrating plasmids based on the TG1 and R4 recombination systems successfully integrated into the expected *attB* sites in A. *marina* DSM45569. Conservation between the *attB* sites for TG1 and R4 in a number of *Amycolatopsis* species is high, suggesting that plasmids with the integration systems from these phages should be widely useful in this genus.

As is common with serine integrase-mediated recombination, the *attB* sites in A. *marina* are located within open reading frames and potentially disrupt the gene. The TG1 *attB^Am^* site is located within a gene predicted to encode a putative succinyldiaminopimelate transaminase (WP_091671332.1), and the R4 *attB^Am^* site is located within a gene predicted to encode a fatty-acyl-CoA synthase (SFB62308.1). Compared to the wild-type (unintegrated) strain, the strains with integrated pHG4 or pJH1R4 did not show any difference in growth. However, further study is required to investigate the effects of TG1 or R4 plasmid recombination on both primary and secondary metabolism as, for example, the integration of ϕC31 integrase-based plasmids has been shown to have pleiotropic effects on bacterial physiology(51).

Currently, the two most commonly used methods of bacterial gene transfer are conjugation and electroporation, both of which come with several advantages and disadvantages. Electroporation involves the introduction of pores within bacterial membranes via an electric current to allow mass, unrestricted transfer of genetic material into species(52, 53). The efficiency of transformation, however, is species-dependent(52). Unlike electroporation, conjugation is not limited by the size of vectors that can be transformed, and has been used successfully to transfer entire genomes in *E. coli*(54). Additionally, conjugation uses the transfer of DNA as a single strand that is relatively insensitive to the majority of the restriction systems of the cells(55). Thus conjugation may have advantages over electroporation. In this study, we successfully integrated a plasmid into the *attB* sites for TG1 and R4 integrases by conjugation, thus supplementing the potential gene transfer methods that could be used in the genus *Amycolatopsis*.

In conclusion, we have identified highly conserved sequences of the *attB* sites for TG1 and R4 integrases within the genus *Amycolatopsis* and demonstrated their use in conjugative DNA transfer. The *A. marina* DSM45569 *attB* sites showed slightly lower recombination efficiencies *in vitro* than the previously identified *attB* sites from *Streptomyces* spp. However, this slight reduction is not enough by itself to explain the order of magnitude reductions in conjugation frequencies observed with A. *marina* compared to *Streptomyces* spp. (Table 3). The conjugation frequencies might be increased by optimising conjugation conditions. Alternatively, efficiently used *attB* sites for the widely used vectors, such as those based on phiC31 *int/attP* could be incorporated into the *Amylcolatopsis* genome using TG1 or R4 integrating plasmids as described here. In short, this work shows that integrative vectors are viable and promising tools for the genetic engineering of rare Actinomycetes.

## Acknowledgments

This work was supported by the Biotechnology and Biological Sciences Research Council project grant BB/K003356/1, and Buvani Murugesan acknowledges the receipt of a summer studentship from the Department of Biology, University of York.

